# Production of Rabies virus-specific monoclonal antibodies and evaluation of their neutralizing potential

**DOI:** 10.1101/2022.11.24.517809

**Authors:** Rafik Soliman, Zeinb Hashem, Mahmoud El-Hariri, Heidy Abo Elyazeed, Hassan Aboul-Ella

**Affiliations:** Department of Microbiology, Faculty of Veterinary medicine, Cairo University, Egypt; VACSERA holding Company for vaccine and serum production, Ministry of Health, Egypt

**Keywords:** Rabies virus, monoclonal antibodies (mAbs), Neutralizing potential

## Abstract

Rabies is a severe viral infection that causes acute encephalomyelitis, with a case fatality rate of nearly 100% following the onset of neurological clinical signs. Rabies irreversible clinical signs development can be effectively avoided with post-exposure prophylaxis (PEP), which includes vaccines and anti-rabies immunoglobulins (RIGs); however, there is no treatment for symptomatic rabies. The major PEP protocol faces serious access and implementation obstacles in association with a resource-limited setting, which could be successfully overcome by substituting RIGs for monoclonal antibodies (mAbs). Lower production costs, consistent supply availability, long-term storage/stability, and an improved safety profile are all advantages of mAbs. The current work focuses on the key characteristics of currently developed mAbs against rabies and highlights their potential as a novel therapeutic approach. Using immunizing Freund’s adjuvanted emulsions of inactivated purified Vero cell rabies vaccine (PVRV, VERORAB) produced by *Aventis Pasteur* to immunize the BALB/c mice. The immunized BALB/c mice were tested for the production of anti-rabies virus-specific antibodies using Enzyme-linked immunosorbent assay (ELISA). High-responder mice were selected for the fusion process. Hybridomas recovered from the fusion process were selected and separated from the unfused cells and unfavorable fused cells by using the selective HAT medium. Twelve days post fusion the produced hybrids were screened for production of Rabies virus-specific antibodies using ELISA. Four murine hybridomas secreting rabies virus-specific monoclonal antibodies (mAbs) have been properly developed. These 4 stable hybrids were successfully cloned into 4 stable clones, namely, 1E4, 1E9, 2F3, 4E1. The rabies virus specific monoclonal antibodies produced by the 4 selected hybridomas were of IgM isotype. Using Western Blot technique, the specificity of the produced hybrids was confirmed. The neutralizing potential of the prepared mAbs was evaluated and the efficacy of mAbs cocktail prepared from the 4 hybridomas to protect mice in post exposure therapy was determined. The mAbs cocktail given to mice at 24 hours post infection was able to offer 100% protection to mice challenged with 1000 LD50 of rabies virus strain whereas all control mice developed the disease.

## Introduction

Rabies disease is an acute viral infection of the central nervous system caused by Rabies virus. This disease remains an important public health problem and has a long history that dates back to antiquity **(Wilkinson, 2002, Chalchisa Buzayehu Barecha, *et al*., 2017, Rajendra Singh, *et al*., 2017, Fooks et al., 2017 and WHO, 2021)**. This disease is an acute rapidly fatal neurological disease and to date therapeutic efforts in humans have proved futile except in rare cases in which rabies vaccine was administered prior to the onset of clinical disease **(Jackson, 2003 and Dong-Kun Yang *et al*., 2013)**.

The Rabies virus (RABV) is a single-stranded RNA virus of the genus Lyssavirus, family Rhabdoviridae. It invades the central nervous system, causing acute and zoonotic disease worldwide. The virus genomic RNA encodes five proteins: nucleoprotein (N), phosphoprotein (P), matrix protein (M), glycoprotein (G), and polymerase protein (L). The Glycoproteins are the major surface antigens to which neutralizing antibodies are bound. The Glycoprotein antigen binds to acetylcholine receptors, leading to neurotransmitter release disorder and causing neurological symptoms **(Yousaf *et al*., 2012 and Fanli Yang *et al*., 2020)**.

The Rabies virus is transmitted by the saliva of biting infected mammals. Human rabies remains an important disease in the world and rabies mortality ranks about eleventh of all infectious disease, although it is almost always preventable **(Haupt, 1999, Rajendra *Singh et al*., 2017)**. This virus is normally transmitted from dog to dog with virus-laden saliva of the diseased animal via bite wounds. Non-bite transmission (ingestion, inhalation, and other mucosal exposure) has been occasionally implicated. It has been suggested that infection may result from mouth-licking and through consumption of regurgitated food, both are common practices in many social candidates. Such forms of transmission may have occurred in outbreaks **(Kat *et al*., 1996)**.

The infection of the brain in rabies results in behavioral changes, likely due to infection of neurons in limbic areas, which facilitates transmission by biting in rabies vectors. There is spread of Rabies virus away from the CNS (centrifugal spread) along neuronal pathways, particularly involving the parasympathetic nervous system, which are responsible for infection of the salivary glands, skin, heart, and a variety of other organs. Infectious Rabies virus is secreted in the saliva of rabies vectors, which is important for transmission to other hosts **(Jackson *et al*., 1999 and Bernhard Dietzschold *et al*., 2008 and Rajendra Singh *et al*., 2017, WHO, 2018)**.

Lethal rabies virus infection could be prevented by post exposure prophylaxis (PEP) through the combined administration of a rabies vaccine and rabies immunoglobulin (RIG). Two types of RIGS are used: human RIG (HRIG) and equine RIG (ERIG), both derived from pooled sera of human donors and horses vaccinated against rabies, respectively. The need to replace theses hyperimmune serum preparations is widely recognized, and monoclonal antibodies (mAbs) that neutralize rabies virus (RV) offer the opportunity to do so **(World Health Organization, 2002, Quiambao *et al*., 2008 and Sanne Terryn *et al.,* 2016 and Guilherme Diasde Melo *et al*., 2022)**.

The first monoclonal antibody of predefined specificity was made by fusing the right combination of cells, a spleen cell with the genetic information to produce specific antibody, and a selectable myeloma cell that was adapted to continuous and vigorous growth in culture **(Kohler and Milstein, 1975). The** monoclonal antibodies have the obvious advantages of single specificity and have been produced out of a need for homogenous antibodies as reproducible reagents. Polyclonal antibodies are a minor component in a complex mixture of serum proteins and are a heterogonous mixture of molecules with a wide range of binding affinities **(Hay and Westwood, 2002)**.

Mouse mAbs, as well as human mAbs, have been shown to protect rodents from a lethal rabies virus (RV) challenge **(Schumacher *et al*., 1989, Enssle *et al*., 1991, Hanlon *et al*., 2001, Hanlon *et al*., 2002, Prosniak *et al*., 2003 and Guilherme Diasde Melo *et al*., 2022)**. One of the most potent human mAbs, SO57, which manifests strong activity against a variety of Rabies virus strains, was described by **Dietzschold *et al*., (1990)**.

Experiments carried out at the Wistar institute showed that neutralizing antibodies were most effective in the early stages of infection, and before invasion of the nervous system. Once the nervous system has been invaded, the efficacy of the mAbs is questionable. Rabies mAbs have clear advantages over the products now in use. They are highly specific, their production can be closely controlled, and their quality can be easily monitored.

Since one antibody alone could not be used, because of the possibility of escape variants, the strategy followed by Wistar’s scientists and recommended by WHO was to choose MAbs with different specificities and to pool them into a “cocktail” **(World Health Organization, 1990)**. The mAbs have demonstrated their activity in certain animal models and with the progress of technology their potential ease of production in large quantities at low cost and ease of quality control compared to polyclonal serum is attractive **(World Health Organization, 2002)**.

The aim of the present work was the preparation of Rabies virus-specific monoclonal antibodies, its characterization and evaluation of its neutralizing potentials.

## Material and Methods

### Materials for production of Rabies virus-specific monoclonal antibodies

– **BALB/c mice**: Ten BALB/c mice from the experimental Animals Farm, *VACSERA* company – Ministry of Health. These mice were healthy and were kept under good environmental and nutritional conditions. The BALB/c mice were divided into 2 groups, the first one contains Eight-week-old BALB/c mice that were used for immunization with the Rabies virus vaccine for production of highly immunized splenocytes. The second group composed of 2 adult mice that were used for preparation of peritoneal macrophages that were used as feeder cells.
– **Inactivated rabies virus vaccine “Verorab”**: It is a freeze-dried purified Rabies virus vaccine PVRV, Verorab^®^, WISTAR strain RABIES PM/WI 38-1503-3M grown on Vero cells produced by Aventis Pasteur.
– **Myeloma cell line (Plasmacytoma)**: The P_3_-NS1 cell line was used. These myeloma cells synthesize but does not secrete K light chain. It was kindly provided by the Central Laboratory for Monoclonal Antibody Production (CLMAP)-VACSERA.
– **Antibody enzyme conjugates for ELISA (Sigma): Peroxidase conjugated Anti-mouse polyvalent immunoglobulins** (IgG, IgA, IgM) the affinity isolated antigen specific antibodies developed in goats using purified mouse IgG, IgA, and IgM as immunogens. **Alkaline phosphatase conjugated Anti-mouse IgG** (whole molecule) antibody developed in goat. The isolated antigen specific antibody was adsorbed with rat serum proteins and stored at 2-8 °C.
– **Enzyme substrate for Horse radish peroxidase (Sigma)**: (ABTS): 2, 2 azino-bis (3-ethyl benzthiazoline-6-sulphonic acid). Green color developed when the enzyme reacted with its substrate.
– **Adjuvants**: Two types were used, namely, **complete Freund’s adjuvant (CFA)** (Sigma) that was used only for primary immunization and **incomplete Freund’s adjuvant (IFA)** (Sigma) that was used for booster immunization.

### Materials used for evaluation of the neutralizing and protective potential of rabies-specific mAbs

– **Twelve 6-weeks-old Swiss albino mice (10-12gms**) obtained from Vacsera Laboratory animals farm, Vacsera Company-Ministry of health. These mice were used for the challenge experiment.
– **Rabies virus strain** supplied from Rabies Research Department in Vacsera Company-Ministry of health. This strain was isolated previously in Vacsera laboratory from brains of a rabid dogs. This strain was stored as infected mouse brains at −75°C after titration by mice inoculation.

### Other materials

– **Polyethylene glycol (PEG 1500& 50%) (Sigma)**: Sterile-filtered, endotoxin, and hybridoma tested. It is used as a fusogenic material in fusion between splenocytes and myeloma to give hybrids.
– **Hypoxanthine-Aminopterin-Thymidine (HAT) Medium Supplement 50X (Sigma)**: Selective medium for hybridomas (HAT medium supplement 50X) lyophilized, γ-irradiated powder for use in aseptic procedures and cell culture tested and stored dry at – 20 °C.
– **Hypoxanthine-Thymidine (HT) Medium Supplement 50X (Sigma)**: used as transition medium for HAT-selected hybridomas before transfer to RPMI medium. (HT medium supplement 50X) lyophilized, γ-irradiated powder for use in aseptic procedures and cell culture tested stored dry at −20 °C.
– **OPI Medium Supplement**: Lyophilized, γ-irradiated powder for use in aseptic procedures and cell culture tested, endotoxin free. Used in the preparation of cloning media with HT medium supplement. Stored dry at −20 °C.
– **Dimethyl sulphoxide (DMSO) Research Grade (SERVA)**: It is used as a freezing medium for cells (sterile filtered, endotoxin and hybridoma tested).

## Methods

### Preparation of a stable emulsion of purified Rabies virus vaccine (PVRV-Verorab) in Freund’s adjuvants

An emulsion was prepared from Rabies virus vaccinal antigen and equal volume of Freund’s adjuvant. Stability test was done by dropping the emulsion into a beaker containing water. Stable emulsion was an oil in water emulsion, which would not disperse when dropped into water.

### The immunization schedule for production of immune spleen cells

Six successive doses of the Rabies virus vaccine **(PVRV-Verorab)** antigens were used. Each dose was injected in the mice according to the following scheme:
– **First immunization of BALB/c mice**: An emulsion containing equal volumes of Rabies virus vaccinal antigens (35μg of the rabies vaccine proteins) and complete Freund’s adjuvant was injected I/P in mice using a dose of 0.3 ml/mouse.
– **Second immunization of BALB/c mice**: Two weeks after the first immunization dose, an emulsion containing equal volumes of Rabies virus vaccinal antigen emulsified in incomplete Freund’s adjuvant was injected I/P in the immunized mice using 0.3 ml/mouse.
– **Third, fourth and fifth immunization of BALB/c mice**: These booster doses were done exactly like the second dose at 2 weeks interval.
– **Final booster dose**: Four days before fusion, 0.3 ml of the Rabies virus vaccine alone was injected I/P.

### Preparation of myeloma cells for fusion with splenocytes from immunized mice according to Ausubel et al., (1990) and Ali (2001)

The myeloma cells were maintained in 250ml tissue culture flasks in 50ml of complete RPMI-1640 culture medium with 20% FCS under environmental conditions of 5% CO_2_, 37°C and 98% relative humidity incubator. The density of the cells was between 10^7^-10^8^ cells/ml. Prior to fusion, the P3-NS1 myeloma cells were in the log phase of growth (*i.e*., over 95% of cells were viable).

### Preparation of splenocytes from immunized BALB/C mice

Briefly, four days after the last immunization, the titer of anti-rabies virus specific antibodies was measured by ELISA and the high responder mouse was selected. The selected mouse was scarified, the spleen was removed, minced and the splenocytes were harvested.

### Polyethylene glycol aided fusion (Ali, 2001)

The splenocytes were resuspended in RPMI supplemented with 20%FCS. The tube was then centrifuged at 1500 for 5min and resuspended in 5ml RPMI supplemented with 20% FCS. The harvested spleen cells and the myeloma cells were counted and adjusted to a ratio of 10:1, respectively. The spleen cells were added to myeloma in centrifugation tube then pelleted together. The Pelleted cells were washed with 10ml serum free medium in centrifugation tube and the supernatant was discarded. To the packed cells 1ml 50% PEG 1500 was added drop by drop over 1 min with tube rotation in the hand. The tube was then put in water bath at 37 °C for 1 min. with gentle shaking every now and then. After centrifugation at 100xg, which is equal to 700 rpm for1 min at 37 °C, 1ml RPMI medium without serum was added drop by drop over 1 min with gentle agitation. Another 4.5 ml RPMI medium without serum were added drop by drop over 3 min with gentle agitation. Five ml RPMI medium without serum were added over 2min with gentle agitation. Then RPMI medium without serum were added to reach a total volume 50 ml. This was followed by centrifugation at 200xg for 5min at 37 °C, the supernatant was discarded, and the cell pellet was gently resuspended in 9ml complete RPMI culture medium. The cell mixture was distributed in three 4 Tissue culture plates (240 well). The outside wells of plates were filled only with RPMI medium. In the first tissue culture plate, the cell suspension was distributed in 100μl/well). Six ml complete RPMI culture medium were added to the cell suspension, mixed, and distributed in the second plate. For the third plate, 3 ml complete RPMI culture medium were added and the cells were mixed and distributed in fourth plate. The plates were left in 5% CO2, 37°C incubator overnight before adding HAT medium.

### HAT selection (Shivnand 2010)

The 96-well plates were changed to HAT medium on the second day following fusion. Half the old medium was removed and replaced with HAT medium (2 ml HAT + 48 ml RPMI) (2X). This medium was changed by HAT medium (1ml HAT + 49 ml RPMI) (1X) on day 4, 6, 8, and every two to three days afterwards until the first screen. Within three to four days all the myeloma cells died and from about the fifth day hybrids were started to become visible. Specific antibody production (at day 12-14 day) was screened by ELISA, at least three days were allowed between screen and last medium change. After screening, positive antibody secreting wells were selected and expanded.

### Expansion of culture prior to cloning. Treatment of the cells with HT medium before cloning (Shivnand 2010)

Using 24 wells tissue culture plate 1ml HT medium was added to each 2ml well. The cells in fusion plate-positive wells were suspended and 100μl/well was transferred to 2ml well and mixed, then 100μl of this cell mixture was returned to the original well in the fusion plate. Two days later much supernatant was removed as possible and fresh HT medium was added. When cells were nearly confluent (up to about one week) the new wells and the original wells were re-assayed for rabies specific antibodies using ELISA. The Hybridoma cells secreting Rabies virus specific mAbs were then cloned by limiting dilution, propagated, and stored in liquid nitrogen.

### Characterization of the positive hybridoma producing Rabies virus-specific monoclonal antibodies

The class of the produced specific Mab was determined using agar gel precipitation test and the mouse immunoglobulin isotyping kits **(Mouse Type Isotyping kit, BioRad, Hercules, CA, USA**) and its specificity was determined using Western blot test according to **Bolt and Mahoney (1997)**.

### Evaluation of the neutralizing potential of the mAbs

The protecting capability and neutralizing potential of the anti-rabies mAbs cocktail prepared from the 4 recovered hybridomas was determined in mice challenge experiment with Rabies virus strain supplied from rabies research Department in Vacsera Company-Ministry of health. This strain was isolated previously in Vacsera laboratory from brains of a rabid dogs. This strain was stored as infected mouse brains at −75°C after titration by mice inoculation.

Twelve 6-week-old Swiss albino mice (10-12gms) obtained from Vacsera Laboratory animals farm, Vacsera Company-Ministry of health, were used for the challenge experiments. In the challenge experiments 1000 LD50 of the Rabies virus strain prepared as 10% mouse brain homogenate in sterile phosphate buffered saline (PBS) was prepared and used as follow: A group of six mice were inoculated intramuscularly in the hind limb with 0.1 ml of virus suspension containing 1000 LD50 of virus. The mice were then inoculated with 1.5 IU/0.1 ml of the mAbs cocktail at the same site after 24 hours post infection. Another group of six mice were inoculated similarly with the rabies virus followed by injection of 0.1 ml of sterile saline at similar time points following infection. The dose of Mab was calculated based on the presently advocated dose of ERIG for passive immunization of humans (40IU/ kg body weight).

## Results

The total protein content of Rabies virus vaccine used for immunization of BALAB/c mice was 8 mg/ml as measured with Biuret test.

### Monitoring of immunization of BALB/C mice with rabies vaccine

The group of eight immunized mice was bled two weeks after the third immunization, and serum was collected. Using ELISA, the developed anti-rabies antibodies titer developed against rabies vaccine was measured. Although several mice manifested good immune response, mice number 6 was selected for the fusion experiment, because of its higher response **(Table 1)**.

**Tab. 1:** Measurement of anti-rabies polyclonal antibodies in serum of BALB/c mice immunized with rabies vaccine using ELISA.

| Immunized mice | Titer of anti-rabies antibodies |  |  |  |  |  |  |  |
| --- | --- | --- | --- | --- | --- | --- | --- | --- |
|  | 1/10 | 1/20 | 1/40 | 1/80 | 1/160 | 1/320 | 1/640 | 1/1280 |
|  | Results were expressed as optical density reading OD |  |  |  |  |  |  |  |
| Negative control | 0.511 | 0.500 | 0.443 | 0.296 | 0.281 | 0.144 | 0.106 | 0.063 |
| Mice 1 | 1.717<br>+ | 1.434<br>+ | 1.082<br>+ | 0.731<br>+ | 0.391<br>+ | 0.114<br>- | 0.058<br>- | 0.039<br>- |
| Mice 2 | 1.801<br>+ | 1.466<br>+ | 1.132<br>+ | 0.772<br>+ | 0.402<br>+ | 0.149<br>± | 0.106<br>- | 0.001<br>- |
| Mice 3 | 1.737<br>+ | 1.432<br>+ | 1.088<br>+ | 0.696<br>+ | 0.396<br>+ | 0.120<br>- | 0.094<br>- | 0.010<br>- |
| Mice 4 | 1.622<br>+ | 1.338<br>+ | 1.045<br>+ | 0.682<br>+ | 0.369<br>+ | 0.097<br>- | 0.044<br>- | 0.056<br>- |
| Mice 5 | 0.651<br>+ | 0.616<br>+ | 0.588<br>+ | 0.569<br>+ | 0.544<br>+ | 0.439<br>+ | 0.365<br>+ | 0.257<br>+ |
| Mice 6 | 1.663<br>+ | 1.602<br>+ | 1.582<br>+ | 1.580<br>+ | 1.570<br>+ | 1.440<br>+ | 1.395<br>+ | 1.218<br>+ |
| Mice 7 | 0.559<br>+ | 0.511<br>+ | 0.480<br>+ | 0.431<br>+ | 0.354<br>+ | 0.287<br>+ | 0.191<br>+ | 0.017<br>- |
| Mice 8 | 1.162<br>+ | 1.160<br>+ | 1.139<br>+ | 1.074<br>+ | 0.975<br>+ | 0.884<br>+ | 0.695<br>+ | 0.451<br>+ |

### Results of fusion of myeloma cells with rabies-immunized spleen lymphocytes

#### Microscopical examination of the fusion plates

Twelve days post fusion, the three fusion plates (240 wells) were screened by microscopical examination. Hybrid cell growth covering 10% to 50% of the surface area of the wells was selected. From the examined 240 wells (four plates) only 76 wells (32%) showed hybrid growth **(Table 2)**.

**Table 2:** Results of microscopical examination of fusion plates 12 days post fusion:

| Plate No. | Total wells No. | Wells showing growth |  | Wells showing dead cells |  |
| --- | --- | --- | --- | --- | --- |
|  |  | No. | % | No. | % |
| 1 | 60 | 41 | 68 | 19 | 32 |
| 2 | 60 | 17 | 28 | 43 | 72 |
| 3 | 60 | 12 | 20 | 48 | 80 |
| 4 | 60 | 6 | 10 | 54 | 90 |
| Total | 240 | 76 | 32 | 164 | 68 |

#### Examination of the tissue culture supernatants from wells showing hybrid growth for anti-rabies antibodies

The first screening for anti-rabies antibody production using ELISA was made 12 days post fusion on supernatants from the hybrid cell growth that covers 10% to 50% of the surface area of the wells. In these wells changing of the medium was stopped 5 days before screening. Only 14 wells (5.8 %) from the total 240 wells were positive **(Table 3)**.

**Table (3).**
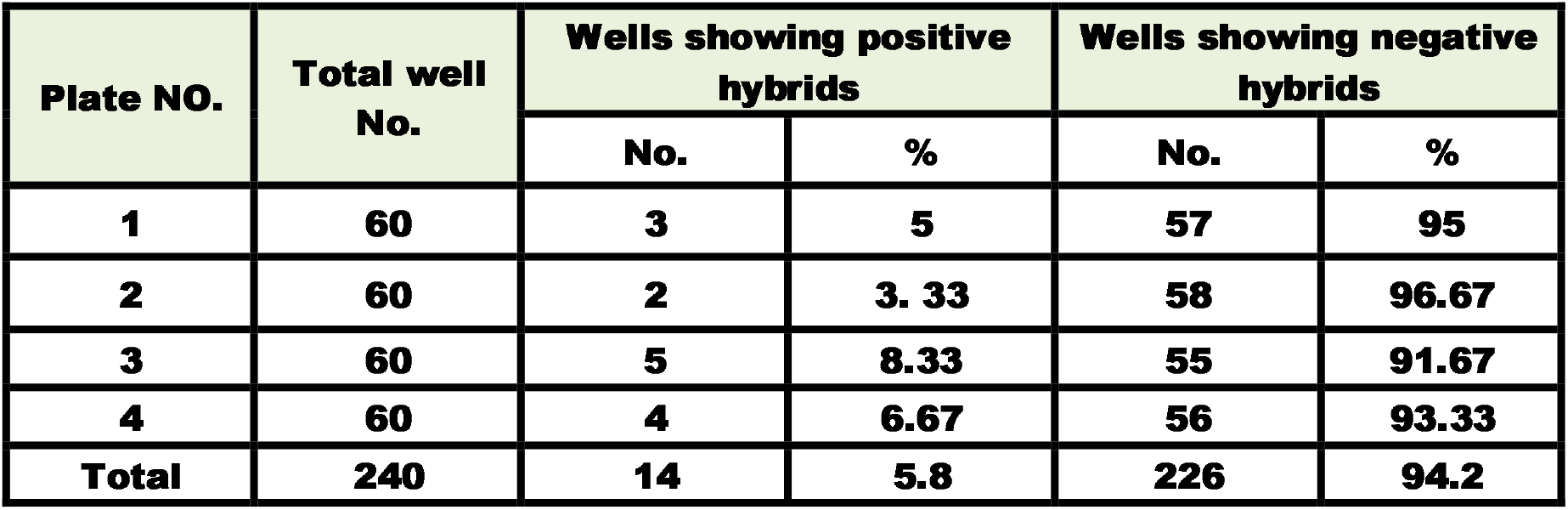
Results of screening of the fusion plate for anti-rabies antibodies producing hybridoma using ELISA 12 days post fusion.

#### Number of stable hybrids producing rabies virus specific monoclonal antibodies

The positive wells were selected and transferred to 24 well plates to grow the cells further. Only four hybrids showed stability after propagation in 24 well plates and the other 10 hybrids did not survive. The positive stable hybrids were specified with its location.

#### Results of cloning of positive hybridomas using limiting dilution method

Cloning of the recovered positive hybridomas using the limiting dilution technique was done to select a pure hybrid cells-producing monoclonal antibodies against rabies virus. The main aim of cloning was to purify the positive hybridoma from non-producer cells. The Cloning plates were examined microscopically and serologically using ELISA to select wells having single clones that produce anti-rabies monoclonal antibodies.

#### Results of screening of the cloned hybridomas using ELISA

Using indirect ELISA revealed that the hybridomas were successfully cloned and four stable clones producing Rabies virus specific mAbs were recovered, namely, **1E4, 1E9, 2F3, 4E1**. A confirmatory testing of the clones’ supernatant using Western Blot test showed positive results of the four obtained clones.

#### Isotyping of the monoclonal antibodies produced by the four clones

Using agar gel procedure and mouse immunoglobulin isotyping kits, all the four obtained clones have been identified and all were found to belong to IgM isotype.

#### The neutralizing potential and protective efficacy of mAbs cocktail prepared from the 4 developed hybridomas

The mAbs cocktail prepared from the 4 hybridomas was able to offer 100% protection to mice challenged with 1000 LD50 of CVS whereas all challenged control mice developed the disease.

## Discussion

Rabies is an acute viral infection of the central nervous system caused by rabies virus, which is transmitted by the saliva of wide range of warm blooded infected mammals. Human rabies remains an important disease in the world and rabies mortality ranks about eleventh of all infectious disease, although it is almost always preventable. Rabies virus was described as a rod- or bullet-shaped, single-stranded, negative-sense, unsegmented, and enveloped RNA virus, measuring approximately 60nm x 180nm. It is composed of an internal protein core or nucleocapsid, containing the nucleic acid, and an outer envelope, a lipid-containing bilayer covered with transmembrane glycoprotein spikes **(Thomas Moulenat *et al*., 2020)**.

The infection of the brain in rabies results in behavioral changes, likely due to infection of neurons in limbic areas, which facilitates transmission by biting in rabies vectors. There is spread of rabies virus away from the CNS (centrifugal spread) along neuronal pathways, particularly involving the parasympathetic nervous system, which is responsible for infection of the salivary glands, skin, heart, and a variety of other organs. Infectious rabies virus is secreted in the saliva of rabies vectors, which is important for transmission to other hosts (**Dietzschold *et al*., 2005)**.

Lethal rabies is prevented by post exposure prophylaxis (PEP) through the combined administration of a rabies vaccine and rabies immune globulin (RIG). Two types of RIGS are used: human RIG (HRIG) and equine RIG (ERIG), both derived from pooled sera of human donors and horses vaccinated against rabies vaccine, respectively **(Omesh Kumar Bharti et al., 2017)**. The need to replace these hyperimmune serum preparations is widely recognized, and monoclonal antibodies (mAbs) that neutralize rabies virus (RV) offer the opportunity to do so.

In 1975, **Kohler and Milstein** made the first monoclonal antibody of predefined specificity by fusing the right combination of cells, a spleen cell with the genetic information to produce specific antibody, and a selectable myeloma cell that was adapted to continuous and vigorous growth in culture. Although, polyclonal antibodies have been used with success for century for therapeutic and prophylactic treatment of infectious diseases, monoclonal antibodies (mAbs) are replacing now the polyclonal antisera production offering substantial advantages in terms of specificity, potency, reproducibility, and freedom from contaminants **(Daniela A. Quinteros, *et al*., 2017)**. On the other hand, the mAbs market has experienced an explosive growth in the last 30 years and more than 200 mAbs have been approved for therapeutic purposes and several hundred are approved for diagnostic approach making this technology i.e., the monoclonal antibody production, one of the fastest growing fields of biotechnology and biopharmaceutical industry and it is estimated that this technology will represents more than 60% of the total biopharmaceutical industry by 2030.

The production of monoclonal antibodies against rabies virus has many advantages over the polyclonal antibodies (serum). It is highly specific and have a low tendency to produce hypersensitivity reaction that can be caused by the polyclonal antibodies also their production can be closely controlled. The quality of the mAbs can be easily monitored and it only needs lab for production rather than polyclonal antibodies which need farms. Since one antibody alone could not be used, because of the possibility of escape variants, the strategy followed by Wistar’s scientists was to choose MAbs with different specificities and to pool them into a “cocktail”. The present work was planned to produce monoclonal antibodies against rabies virus antigens and to characterize it as a preliminary step toward the preparation of monoclonal therapeutic product.

In the present work, before fusion procedure the tissue culture plates were supplemented with peritoneal macrophages as feeder cells, which support the growth of the hybridoma. This goes hand to hand with that done by **Astaldi *et al*., (1980)**, Although other types of feeder cells can be used, e.g., thymocytes, macrophages as feeder cells probably secrete growth factors, which stimulate the growth of the hybridomas, remove toxic by-product from media and exert a synergistic effect. Also, it increases markedly the yield of hybrids and remove dead parental cells and debris from the cultures **(Quinlan and Kennedy, 1994 and Clark, 1995)**.

In concern with the fusion procedure, the splenocytes (B-cells) harvested from a hyper-immunized BALB/C mice were fused with myeloma cell lines (P3NS1). The myeloma cells were selected in log phase with viability over 90%. The fusion between B-cells and myeloma cells was supported by the polyethylene glycol (PEG) 1500, 40% concentration. PEG is a polywax material that promotes cell adherence and exchange of their nuclei. It induces the formation of somatic cell hybrids and facilitated an improved and simplified technique for cell fusion in vitro **(Davidson and Gerald, 1976)**. Normally cell fusion can occur spontaneously in culture at low levels, but its incidence can be increased by treatment of cells with a fusogenic agent like **PEG (Caroline Macdonald, 1998, Hunt and Zola, 2000, Little *et al*., 2000 and Zola *et al*., 2002)**. This agent still is widely used in infusion technology by most laboratories at a concentration of 40-50% (W/V). In fusion technology different ratios between myeloma cells and splenocytes have been tried by several authors. In the present work 10:1 cell ratio between splenocytes and myeloma has been used. This is similar to what was done by **Kenimer *et al*., (1983)** who fused (10^8^) spleen cells and (10^7^) myeloma cells in fusion process in the presence of 50% PEG 1000 solution added to the mixture of both cells.

The produced spleen-myeloma cells hybridomas, which were selected from the unfused cells and undesirable fused cells, using hypoxanthine aminopterin thymidine (HAT selective medium) according to **Kohler and Milstein (1975); Scheidegger and Groth (1980) and Little *et al*., (2000)**. Exposure of fusion cultures to a selective medium with hypoxanthine, aminopterin and thymidine reduced the labor involved and increased the yield. Cell fusion is random; therefore, the cell culture contains a mixture of myeloma-spleen cells fusion, myeloma-myeloma cells fusion, and spleen-spleen cells fusion. Selection of myeloma-spleen cells fusion can be only accomplished by culturing the cell mixture in hypoxanthine aminopterine thymidine (HAT) medium. A Plasmacytoma cell line is deficient in enzyme hypoxanthine guanine phosphoribosyl transferase (HGPRT) responsible for incorporation of hypoxanthine.

Post fusion care started seven days after plating out the cells in HAT medium with replacement of half the medium with fresh medium containing HT, instead of HAT. At this time, small hybridoma growth was detected at the margin of several wells. Supernatants from these wells were collected and tested for the presence of mAbs using ELISA according to **Hunt and Zola, (2000)**. In the present work, 12 days post fusion, hybridoma cell growth that covers 10% – 50% of the surface area of the wells were screened for antibody production against rabies virus using ELISA. Most of researchers use ELISA for screening of monoclonal antibody production by hybridoma **(Kenimer *et al*., 1983; Hunt and Zola, 2000 and Kane and Banks, 2000)**.

Four positive hybridoma were successfully recovered producing specific monoclonal antibodies against rabies virus. These hybridoma cell lines were cloned successfully. Cloning of the positive hybridoma was done using limiting dilution technique to isolate single stable clones producing specific monoclonal antibodies against rabies virus. Cloning was repeated two times in order to remove the non-producing cells from the clones and to avoid the overlapping of the antibody producing cells by the non-producer ones. Also, to guarantee the monoclonality of the antibody producing cell (i.e., to confirm its derivation from a single hybrid cell). This agrees with that stated by **Kennett, (1979); Kennett, (1980); Mike Clark, (1995) and Little *et al*., (2000)**.

The prepared Rabies virus-specific mAbs cocktail from the 4 cloned hybridomas, which are of IgM isotypes, proved effective in inducing 100% protection against rabies virus infection when given to rabies virus challenged mice 24hrs post challenge as compared with to control challenged non treated mice, which died from rabies challenge with rabies virus **(Guilherme Diasde Melo *et al*., 2022)**.

